# The Effects of Central Nervous System Stimulants on *Drosophila melanogaster* Reproduction

**DOI:** 10.1101/145896

**Authors:** A.S. Blake McMahon

**Affiliations:** Diablo Valley College, Department of Biological Sciences, Pleasant Hill, CA, 94523

## Abstract

Stimulant drugs are used everyday by people around the world. The effect stimulants have on developing human fetuses is widely unknown. The fruit fly *Drosophila melanogaster* has become a valuable system to model the complex effects and properties of drugs in mammals. In this study, *Drosophila* is used to analyze the effects of stimulant exposure on reproduction to determine if stimulants cause a significant decrease in the number of offspring produced by parent generations. Caffeine, nicotine, and pseudoephedrine hydrochloride were found to significantly decrease the number of offspring in experimental populations. Further experimentation is necessary to understand the mechanisms underlying these results.

## Introduction

Stimulants are an unavoidable, invisible driving force in todays society. The coffee, tea, and energy drink industries are booming; everyday the average American consumes 168 mg of caffeine from a variety of sources (Fredholm, *et. al.* 1999). Chronic caffeine intake does not appear to negatively affect health, and caffeine users report easily maintaining control of caffeine intake. The widespread use of caffeine, little or no known negative health effects, and the lack of evidence suggesting addiction is a concern have led to caffeine becoming the most widely consumed psychoactive substance in the world (Daly, 1998).

Nicotine, consumed in the form of tobacco, gum, and e-cigarettes, is also one of the most consumed stimulants in the world. The World Health Organization reports that there are 1.27 billion tobacco users world wide (Unknown Author 2, 2016). The IARC monograph has yet to classify nicotine as a carcinogen, yet a study has found nicotine to be potentially carcinogenic (Mishra, *et. al.* 2015). Nicotine also has severe effects on different body systems and is highly addictive (Mishra, *et. al.* 2015). This has led to global regulation of nicotine.

Prescription pharmaceuticals provide another source of stimulants. The CDC reports that 6.4 million children were diagnosed with ADHD, a disorder that is often treated with prescription stimulant medication (Unknown Author 1, 2016). Adderall^®^, a mixture of amphetamine salts, is one of the most common prescription amphetamines used to treat ADHD. While amphetamine salts and amphetamines in general are safe to use at therapeutic dosages, addiction potential is still very high (Spiller, *et. al.* 2013).

Stimulants are also present in less obvious places; pseudoephedrine, a biological precursor to methamphetamine, is found in many over-the-counter nasal decongestants. The regulation of pseudoephedrine has largely been guided by its use in methamphetamine manufacturing and the war on drugs (Unknown Author 3, 2016).

Stimulants provide immediate, measurable feedback by affecting productivity, memory, attention, and energy levels. However, there is a lack of information available on the long term effects of stimulants on prenatal development. Amphetamine salts, caffeine, and pseudoephedrine are classified in the US Pregnancy category “C class” of drugs, suggesting limited information is available on how or if the stimulants affect developing fetuses, and risks have not been ruled out.

The goal of this study was to begin investigating the effects of these stimulants on the model organism *Drosophila melanogaster*, and to determine whether or not any harmful effects were observed after exposing multiple generations of *Drosophila* to the stimulants. A “harmful effect” for the purpose of this study, was defined as a statistically significant decrease in the number of offspring counted in the experimental vials, as compared to the number of offspring counted in the control vials. Information about the effects of stimulants on *Drosophila* could lead to a greater insight into human health and how our interactions with stimulants today will effect the next generation of adults.

Researchers have found that *Drosophila* has dopaminergic neuroreceptors that respond to CNS stimulants in a similar fashion to humans (Bainton, *et. al.* 2000). Additionally, nicotine has been shown to positively affect flies that have motor coordination problems comparable to the symptoms of Parkinson’s disease in humans (Chambers, *et. al.* 2013). *Drosophila* brains have been used in extensive studies to illustrate the mechanisms of amphetamine action in synaptic vesicles (Freyberg, *et al.* 2016). Lastly, a review of studies that used *Drosophila* as a model organism concluded that the molecular mechanisms found in *Drosophila* which control drug-induced behavior have also been found in mammals, validating the use of this model for drug-related research (Kaun, *et. al.* 2012). While many of these studies focus on how the drugs affect addiction and reward pathways in the brain, some research has also suggested an association between drug exposure and reproduction in *Drosophila*. A study conducted at the University of Nebraska found that chronic amphetamine salt ingestion by *Drosophila* caused progressively higher rates of mutation in succeeding generations relative to controls, resulting in physical deformities related to HOX gene expression. This clearly indicates that stimulants can affect the body of *Drosophila* as well as its nervous system. (Surridge, 1972).

As a result of these studies, *Drosophila* is well accepted as a model organism for predicting the effects of CNS stimulants on the number of offspring produced.

I hypothesized that the original mating group of flies in vials exposed to stimulants will produce fewer flies in subsequent generations relative to control populations. This effect was expected to be more pronounced in vials exposed to higher concentrations of stimulants.

## Materials and methods

*Drosophila melanogaster* was exposed to four CNS stimulants: amphetamine salts, caffeine, nicotine, and pseudoephedrine. Caffeine was purchased in pure powdered form from Carolina Biological Supply. A 4.8% nicotine solution with a vegetable-glycerin base was purchased from Wizard Labs. 30mg pseudoephedrine tablets were purchased over-the-counter at a pharmacy. The source of the amphetamine salts used in this study was Adderall®, a schedule II prescription medication. It was obtained through a private donation. 10mg instant release Adderall® tablets were used.

All substances are soluble in water, so flies were exposed to the stimulants by hydrating the growth medium with aqueous solution. Four concentrations were selected for this study (1.0, 0.1, 0.01, and 0.001 mM). This dosing method was modeled after the results of a pilot study conducted to find concentrations of stimulants that did not prove lethal to the original mating group. Solutions were prepared as follows:

### Caffeine

A 1.0mM solution was prepared by mixing the pure powdered caffeine into deionized water at 60 °C, and stirring until dissolved. Serial dilutions were then made to prepare the 0.1, 0.01, and 0.001 mM solutions.

### Amphetamine Salts

Adderall^®^ was used as the source of amphetamine salts. 20 10mg pills were crushed to a fine powder using a mortar and pestle, and a 1.0mM solution was prepared by mixing the powdered Adderall^®^ into deionized water at room temperature and stirring until dissolved. Serial dilutions were then made to prepare the 0.1, 0.01, and 0.001 mM solutions. It was determined by weight that 23mg of power contained 1mg of amphetamine salts.

### Pseudoephedrine Hydrochloride

30 mg pills form and must be crushed. Using a mortar and pestle crush the pseudoephedrine until it is a fine powder. 5.5 mg of powder contains 1 mg of pseudoephedrine. 20 30mg pills were crushed to a fine powder using a mortar and pestle, and a 1.0mM solution was prepared by mixing the powdered pseudoephedrine hydrochloride into deionized water at room temperature and stirring until dissolved. Serial dilutions were then made to prepare the 0.1, 0.01, and 0.001 mM solutions. It was determined by weight that 5.5mg of power contained 1mg of pseudoephedrine hydrochloride.

### Nicotine

A 4.8% nicotine solution in a vegetable glycerin base was diluted into a 1.0 mM solution using deionized water. Serial dilutions were then made to prepare the 0.1, 0.01, and 0.001 mM solutions.

Adderall^®^ and pseudoephedrine hydrochloride are available in 10 and 30 mg tablets, respectively. These tablets contain fillers, so the ratio of the pure drug to the fillers had to be determined. To do this, 30 tablets of each drug were weighed. Their weight was averaged and a ratio of milligrams of drug to milligrams of drug and filler was calculated. A gram scale sensitive to 1*10^−3^g was used when measuring the dry weight of caffeine, Adderall^®^, and pseudoephedrine. Because nicotine is only available in liquid form, it was measured by volume.

*Drosophila melanogaster* was the species used in this study. *Drosophila* culture vials were purchased from Carolina Biological Supply. A parent generation was bred in the cultures to ensure that all flies placed in the first generation of experimental vials came from the same initial swarm of flies.

In total, 54 *Drosophila* habitat vials were created over three generations, 18 per generation. Each generation had two vials serve as control populations and 16 serve as experimental populations.

The vials were composed of 7.5g of *Drosophila* growth medium, 30mg of yeast for adult flies to eat, 20mL of aqueous-stimulant solution or water depending on a vials classification as experimental or control, respectively, and four male and four female flies. White, undyed *Drosophila* growth medium was purchased from Carolina Biological Supply, yeast was included with this purchase. A volumetric pipet was used to transfer exactly 20.00 mL of solution on to the growth medium in each vial. Each drug was used in four vials, each vial containing a different concentration of drug.

**Figure 1.**
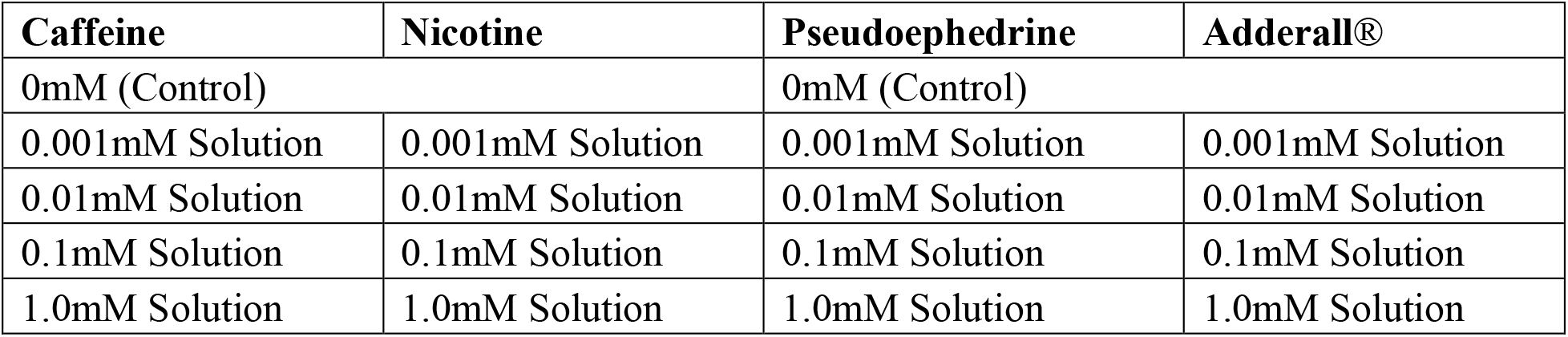
A dosing chart outlining how all 16 experimental vials were dosed. The control vials were hydrated using deionized water. Each drug was used to dose four vials as four different conentrations.

After the vials were created, the culture vials were anesthetized using FlyNap^®^, according to the manufacturers instructions. The anesthesia is a volatile ether compound (Carolina Biological Supply). Flies were examined for gender using a dissecting microscope. Four male, and four female flies were placed in each vial to breed for five days, or 120 hours. The eight adult flies were then anesthetized and removed from the habitat. All vials were kept in an incubator set to 25±1°C. This resulted in a development time (egg to adult) of 10 days. It was crucial that only the 1^st^ generation flies were counted (Parent x Parent cross), so the eggs and pupae were allowed to mature for 9 days after orphaning, ensuring all of the eggs laid by the parent generation were able to mature from the larval stage to the adult stage. This was not enough time for any second generation flies to mature to adulthood. On day 19, 9 days after orphaning, second generation vials were prepared, and the first generation flies were anesthetized. Four male and four female flies were removed from each first generation vial and placed in the second generation vial that was dosed with the same stimulant and concentration used in the their first generation vial. The second generation vials were placed in the incubator.

The first generation vials were placed in the freezer to kill all the flies, and ensure no developing larvae would mature to adulthood, skewing the number of first generation offspring. This method also preserved the adult fruit flies until a vial was ready to be counted. A dissecting microscope was used to count the fruit flies, and identify gender.

The second generation was handled the same way as the first generation, ending in preparing the third generation of vials and freezing the second generation for counting. All three generations of flies assayed were exposed to the same range of stimulant concentrations.

Three generations were assayed so that observations could be made on how chronic exposure to stimulants affects the reproductive capablities of a population over time.

## Results

54 vials contained over three generations and exposed to four different stimulants at four different concentrations produced 23,698 flies, which were counted for analysis. The vials had a consistent male-to-female ratio averaging .848 with a standard deviation ±0.069. Data was first analyzed to determine if the negative effects of stimulant exposure on reproduction would compound in successive experimental generations.

The number of *Drosophila* offspring remained consistent as successive generations were exposed to amphetamine salts; the effect of amphetamine salt exposure on reproduction did not compound generation to generation. Compared to the control population, no concentration of amphetamine salts had a significant effect on the number of offspring present in experimental vials.

The reproductive capabilities of *Drosophila* did change significantly as successive generations were exposed to nicotine. Compared to the control population, nicotine exposure at concentrations of 1.0 and 0.1 mM significantly affected the number of offspring produced, while .01 and .001 mM concentrations had no significant effect.

The negative effects of caffeine exposure compounded in successive experimental generations that were exposed to the varying concentrations of caffeine. Compared to the control population, caffeine exposure at concentrations of 1.0, 0.1, and 0.001 mM significantly affected the number of offspring produced.

The reproductive capabilities of *Drosophila* did change significantly as successive generations were exposed to pseudoephedrine. Compared to the control population, pseudoephedrine exposure at concentrations of 1.0 and 0.1 mM significantly affected the number of offspring produced, while 0.01 and 0.001 mM concentrations had no significant effect.

As subsequent generations of flies were assayed, the mating group placed in the vials became less stable. When the first generation parents were removed, six flies had died over the five day mating period. The second generation parents had 14 dead flies at orphaning, and the third generation had 27 dead flies at orphaning. This decrease in viability was observed in both the experiment and control populations.

**Figure 2.**
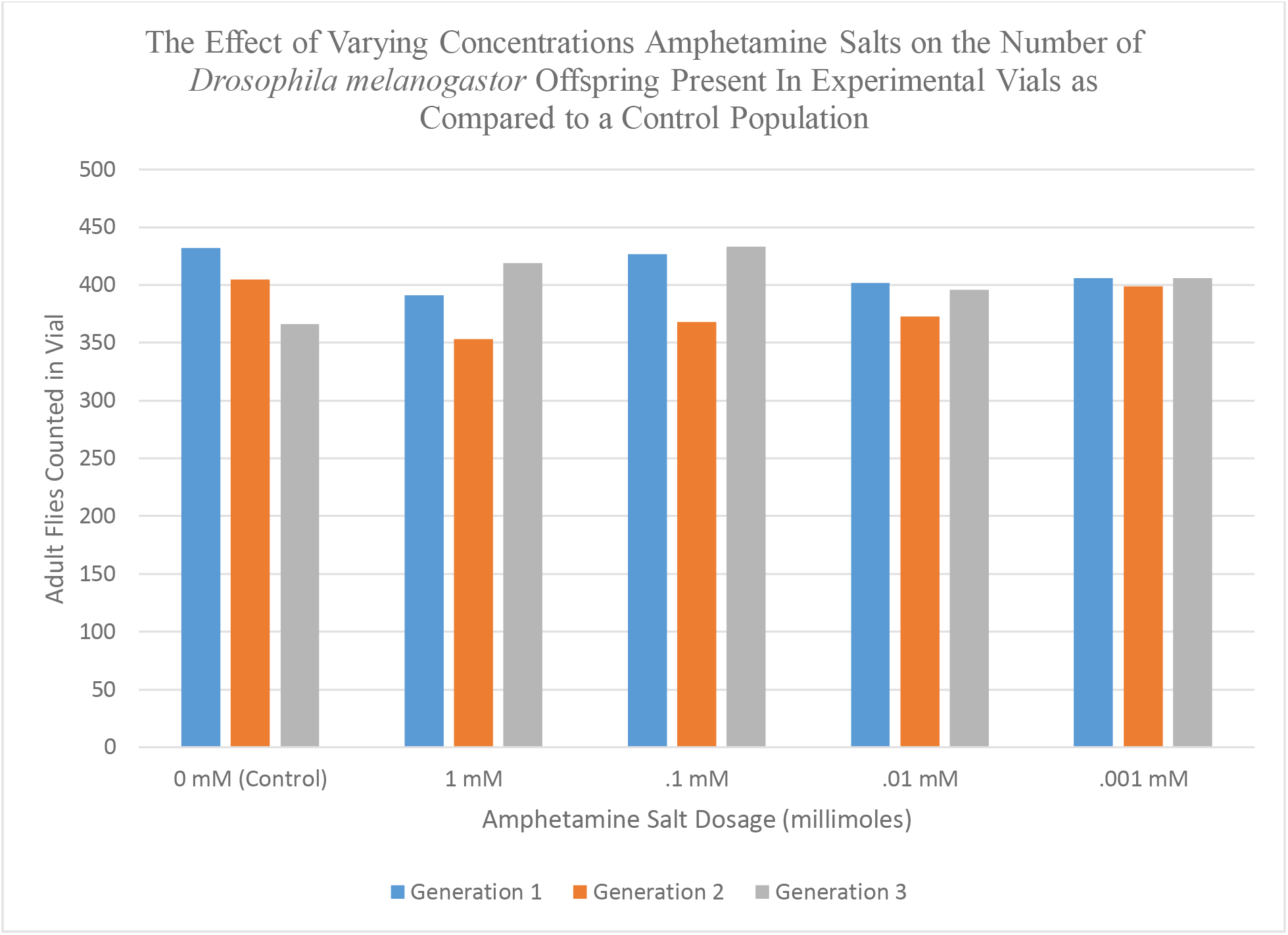
Three generations of *Drosophila melanogaster* were raised on a growth medium hydrated with amphetamine salt in 1, 0.1, 0.01, and 0.001 mM concentrations. The number of offspring from each generation was counted to determine if amphetamine salt had a deleterious effect on the number of offspring present. A students t-test was used to determine that exposure to amphetamine salt showed no significant change in population when compared to the control populations (p-value < 0.05).

**Figure 3.**
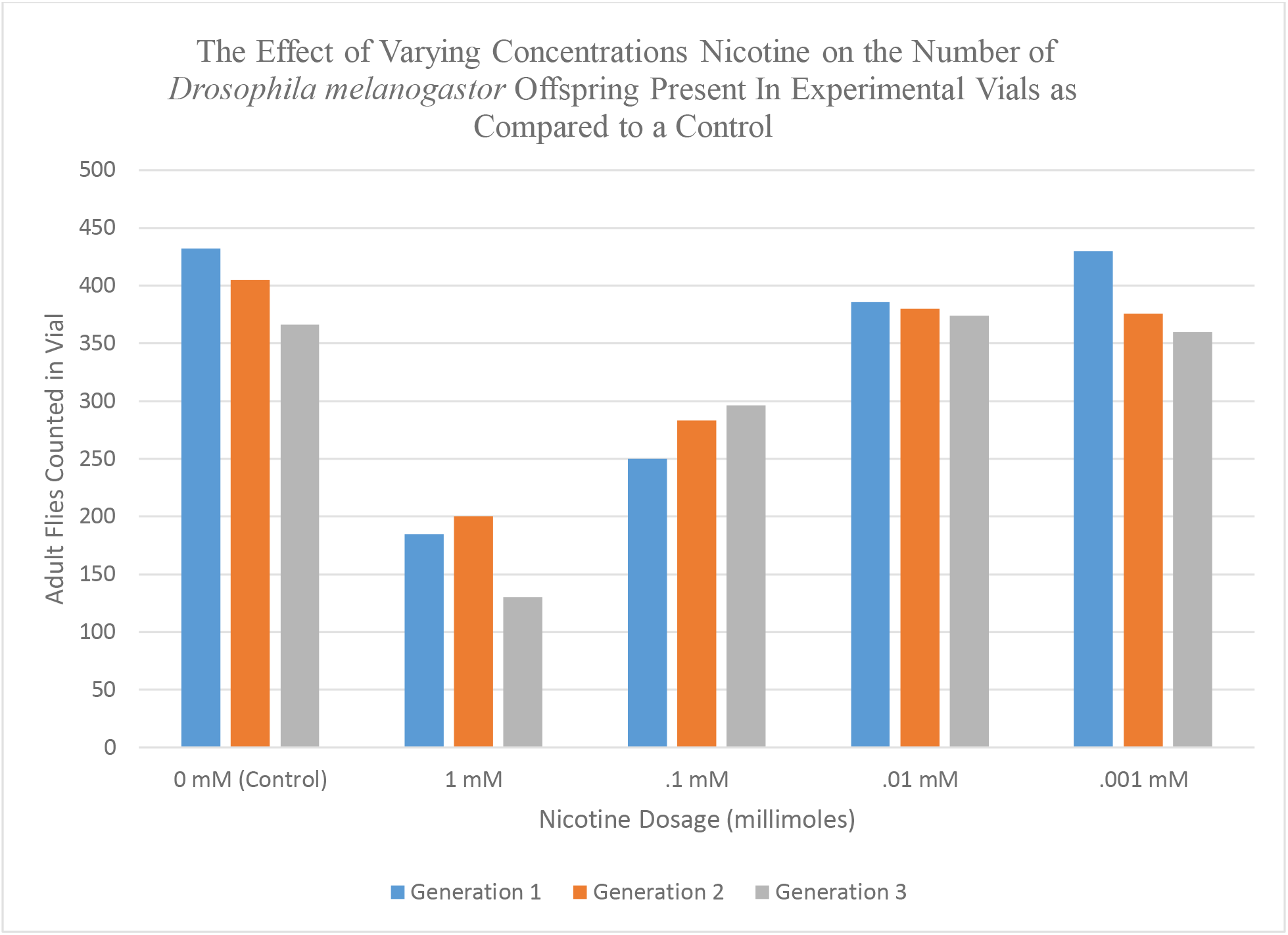
Three generations of *Drosophila melanogaster* were raised on a growth medium hydrated with nicotine in 1, 0.1, 0.01, and 0.001 mM concentrations. The number of offspring from each generation was counted to determine if nicotine had a deleterious effect on the number of offspring present. A students t-test was used to determine that exposure to nicotine caused a significant decrease in the number of offspring produced by populations exposed to 1.0 and 0.1mM concentrations when compared to the control populations (p-value < 0.05).

**Figure 4.**
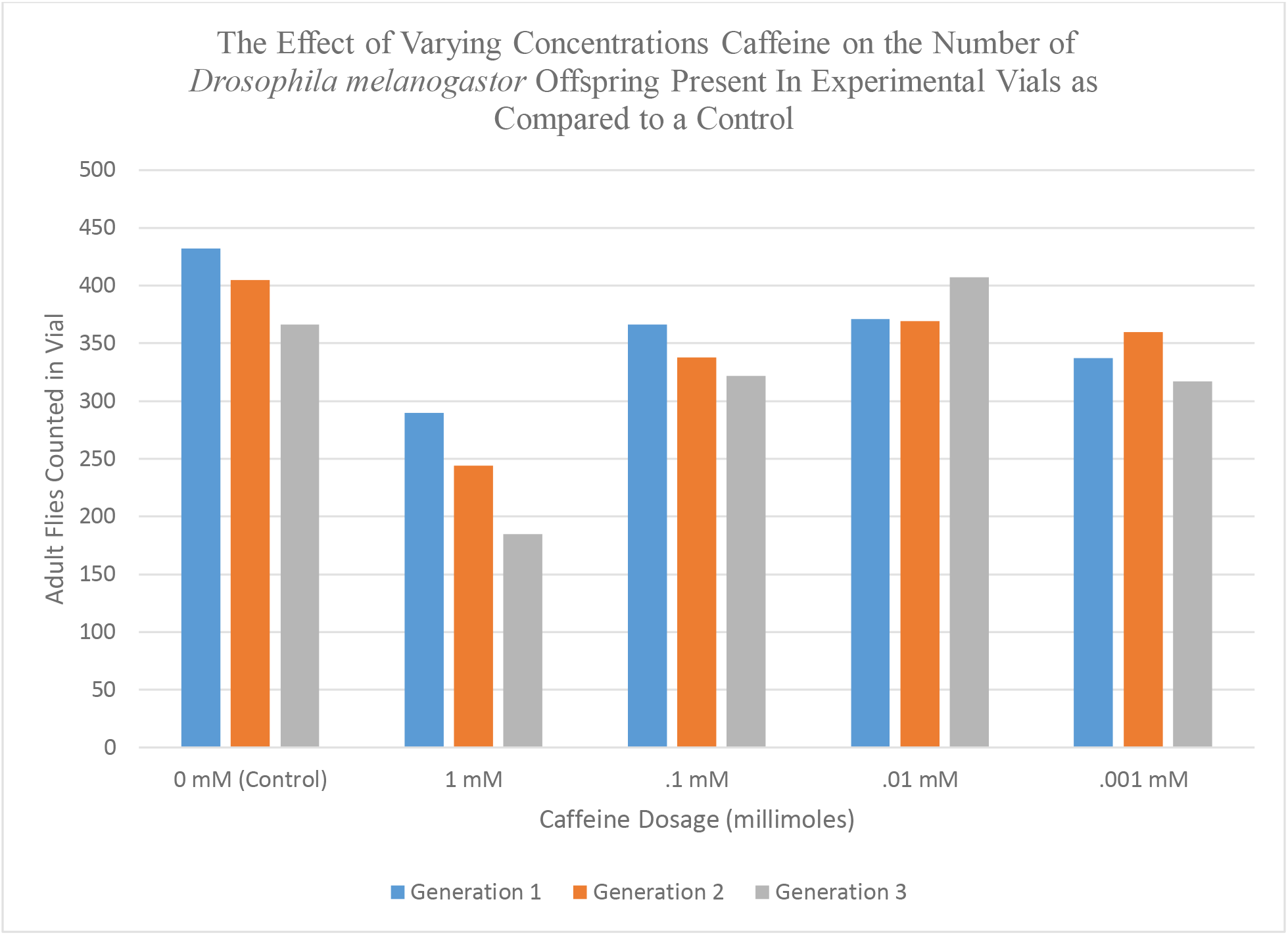
Three generations of *Drosophila melanogaster* were raised on a growth medium hydrated with caffeine in 1, 0.1, 0.01, and 0.001 mM concentrations. The number of offspring from each generation was counted to determine if caffeine had a deleterious effect on the number of offspring present. A students t-test was used to determine that exposure to caffeine caused a significant decrease in the number of offspring produced by populations exposed to 1.0, 0.1, and 0.001mM concentrations when compared to the control populations (p-value < 0.05).

**Figure 5.**
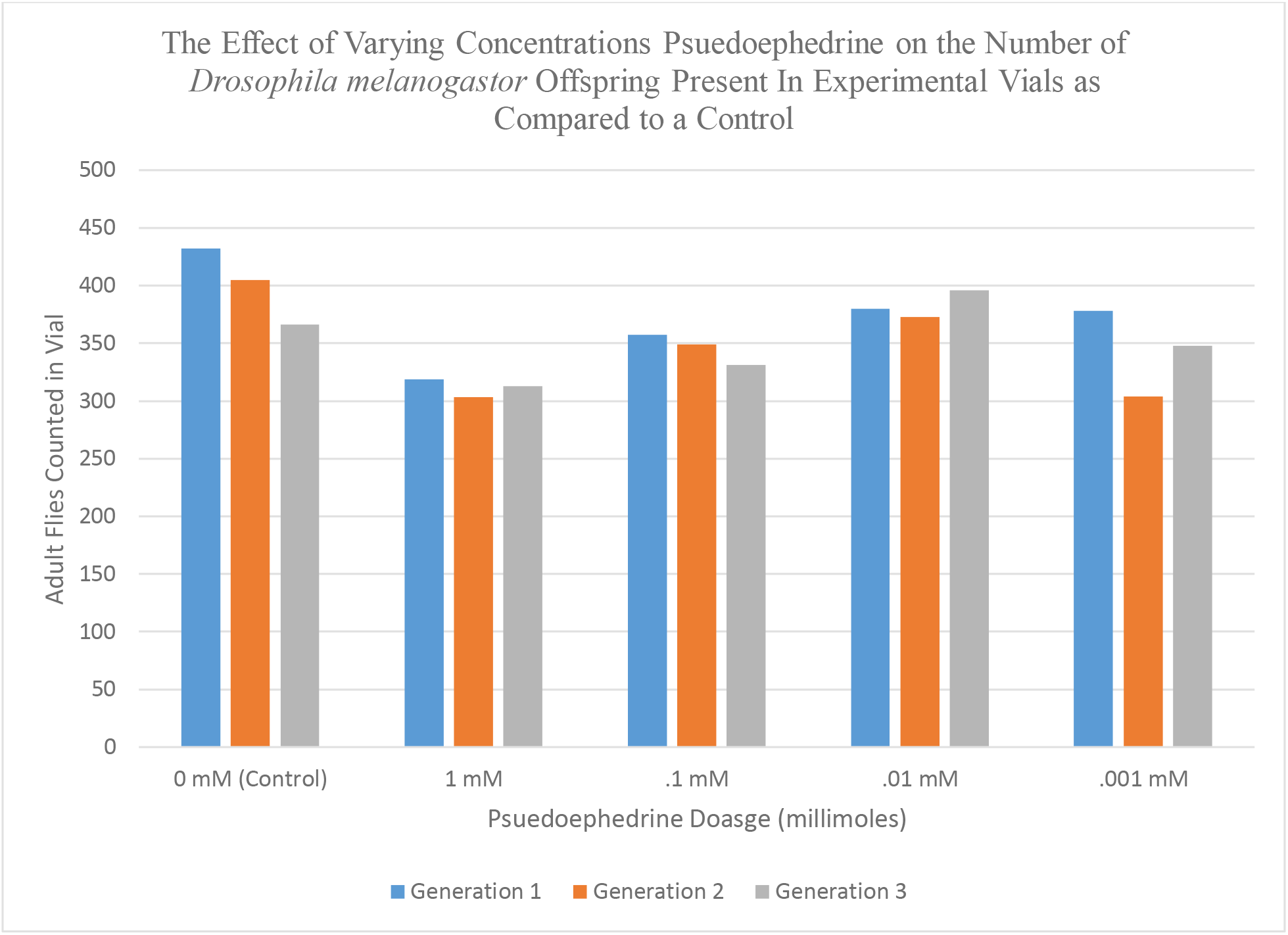
Three generations of *Drosophila melanogaster* were raised on a growth medium hydrated with pseudoephedrine hydrochloride in 1, 0.1, 0.01, and 0.001 mM concentrations. The number of offspring from each generation was counted to determine if pseudoephedrine had a deleterious effect on the number of offspring present. A students t-test was used to determine that exposure to pseudoephedrine caused a significant decrease in the number of offspring produced by populations exposed to 1.0, 0.1, and 0.001mM concentrations when compared to the control populations (p-value < 0.05).

## Discussion

The present experiment offers no explaination to the mechanism underlying the results. Flies were dosed regardless of age, gender, or sexual history. Differentially dosing female and male flies before mating would offer greater insight into how gamete production or embryonic development is affected by exposure. Additionally, counting the number of eggs layed by individual female flies would indicate if fecundity is affected during exposure. The results from other research points to a decrease in sperm viability as a possible explaination for the results seen in this experiment. Primary infertility has been observed in male rats as a result of nicotine ingestion (Oyeyipo, 2011). Nicotine decreased sperm motility and count in the study. A study performed at the University of Massachusetts found that the rate of polyspermic sea urchin eggs was directly proportional to concentrations of nicotine in the environment (Longo, 1970). This study suggests that spermatozoa exposed to nicotine undergo structural changes that make polyspermy more likely, resulting in a decrease in egg viability after fertilization. High caffeine intake (>800mg/day) has been shown to reduce sperm concentration and total sperm count in human men (Jensen, 2010). Pseudoephedrine was found to induce sperm abnormalities, and lower sperm count in rat testis (Thanoi, 2012). These results suggest that the decrease in the number of offspring present in experimental vials of *Drosophila* dosed is due to a decrease in the viability of sperm in male *Drosophila*.

Egg viability should also be tested in further experimentation. If the method of egg collection described by Mary Tyler in <u>Developmental Biology</u> was applied to female flies, fecundity after exposure could be directly measured (Tyler, 2001). This method uses a specialized agar egg laying medium which allows for the exact number of eggs laid to be counted and inspected for viability. Eggs collected by this method should be given time to mature in control and experimental conditions to determine if the viability of fertilized eggs decreases in environments where stimulants are present.

Caffeine and pseudoephedrine were found to significantly decrease the number of offspring in experimental populations dosed with 1, 0.1 and 0.001mM solutions of stimulant but not in the population dosed with a 0.01mM solution (p-value < .05). This result suggests there was an uncontrolled extraneous variable that caused the 0.001mM environments to return significant results incorrectly. Every generation of the pseudoephedrine 0.001mM had at least two of the eight parent flies die during the five day mating period. This problem also affected the second and third generation of the vials exposed to 0.001mM caffeine solutions. However, the vials exposed to 0.01mM pseudoephedrine and caffeine did not experience this problem. Therefore, this result is likely inconsistent because less parent flies survived the mating period in the pseudoephedrine and caffeine 0.001mM vials, so less eggs were laid and less offspring hatched.

